# Differential Effects of endogenous and exogenous attention on sensory tuning

**DOI:** 10.1101/2021.04.03.438325

**Authors:** Antonio Fernández, Sara Okun, Marisa Carrasco

**Affiliations:** Department of Psychology, New York University, New York, NY 10003, USA; Center for Neural Science, New York University, New York, NY 10003, USA

## Abstract

Covert spatial attention (without concurrent eye movements) improves performance in many visual tasks (e.g., orientation discrimination and visual search). However, both covert attention systems, endogenous (voluntary) and exogenous (involuntary), exhibit differential effects on performance in tasks mediated by spatial and temporal resolution suggesting an underlying mechanistic difference. We investigated whether these differences manifest in sensory tuning by assessing whether and how endogenous and exogenous attention differentially alter the representation of two basic visual dimensions: orientation and spatial frequency (SF). The same human observers detected a vertical grating embedded in noise in two separate experiments (with cues to elicit either endogenous or exogenous attention) and reverse correlation was used to infer the underlying neural representation from behavioral responses. Both endogenous and exogenous attention similarly improved performance at the attended location by enhancing the gain of all orientations without changing tuning width. In the SF dimension, endogenous attention enhanced the gain of SFs above and below the target SF, whereas exogenous attention only enhanced those above. Additionally, exogenous attention shifted peak sensitivity to SFs above the target SF, whereas endogenous attention did not. Both covert attention systems modulated sensory tuning via the same computation (gain changes). However, there were differences in the strength of the gain. Compared to endogenous attention, exogenous attention had a stronger orientation gain enhancement but a weaker overall SF gain enhancement. These differences in sensory tuning may underlie differential effects of endogenous and exogenous attention on performance.

## Introduction

Attention improves visual perception by selectively processing incoming information. Such a system is necessary given the high cost of cortical computation and the constant exposure of our visual system to far more information than it can process (Lennie, 2003). There are two types of covert spatial attention: Endogenous–voluntary, goal driven and deployed in ~300ms, and exogenous–involuntary, stimulus driven and deployed fast and transiently, peaking at ~100ms (reviews, Carrasco 2011, 2014). Similar temporal dynamics have been reported from single cell recordings in macaque area MT (Busse et al., 2008). Orienting covert spatial attention to a target location benefits performance in many tasks. These improvements often manifest via an increase in contrast sensitivity—the ability to distinguish an object from its surroundings—and spatial resolution—the ability to see fine detail (reviews, Carrasco 2011, 2014; Anton-Erxleben & Carrasco, 2013; Carrasco & Barbot, 2014).

Differences exist in how endogenous and exogenous attention modulate contrast responses. Endogenous attention typically increases contrast sensitivity via a contrast gain change, whereas exogenous attention increases maximal contrast responses via a response gain change (Ling & Carrasco, 2006; Pestilli et al., 2009; Fernández & Carrasco, 2020; but see Morrone et al., 2002, 2004). A prominent normalization model of attention postulates that these differences are due to changes in the size of the attention field relative to the stimulus size (Reynolds & Heeger, 2009).

Differences also exist in tasks mediated by spatial resolution such as texture segmentation. Endogenous attention optimizes resolution to improve performance at central and peripheral locations (Yeshurun et al., 2008; Barbot & Carrasco, 2017; Jigo & Carrasco, 2018). However, exogenous attention inflexibly increases resolution resulting in performance benefits at peripheral locations and costs at central locations (Yeshurun & Carrasco, 1998, 2000; Talgar & Carrasco, 2002). Adapting observers to high spatial frequencies (SFs) removes the attentional impairment, suggesting that exogenous attention inflexibly increases resolution by enhancing sensitivity to high SFs (Carrasco et al., 2006). Endogenous attention improves performance at all eccentricities by flexibly adjusting resolution via the contribution of high-SFs information according to task demands (Barbot & Carrasco, 2017). Given the tradeoff between spatial and temporal resolution, exogenous attention enhances spatial resolution but degrades temporal resolution (Yeshurun 2004; Yeshurun & Levy, 2003; Rolke et al., 2008) and perceived motion (Yeshurun & Hein, 2011). Conversely, endogenous attention improves both spatial and temporal resolution (Sharp et al., 2018).

A recent image-computable normalization model of attention reproduces the well-established differential effects of endogenous and exogenous attention on contrast sensitivity and spatial resolution by assuming that the attention effects are mediated by differences in SF tuning (Jigo, Heeger & Carrasco, 2021). Here, we investigate, for the first time, whether and how endogenous and exogenous attention differentially mediate sensory representations. We focus on two vision building blocks coded in V1: orientation and SF (Hubel & Wiesel, 1959; Maffei & Florentini, 1977). Sensory representations were assessed using reverse correlation, which is widely used to probe the mechanisms underlying behavioral responses to noisy stimuli (Ahumada, 1996; Eckstein & Ahumada, 2002; Wyart et al., 2012; Li et al., 2016) and can approximate electrophysiological sensory tuning properties (Neri & Levi, 2006). Reverse correlation, which provides a measure of how changes in external noise influence performance, complements conventional methods, like signal detection theory (SDT), that cannot distinguish between external and internal performance limiting noise. Here we used SDT to assess performance with endogenous and exogenous attention and reverse correlation to uncover their underlying computations.

The same observers completed two experiments with identical stimuli and task but different cues. Both endogenous and exogenous attention altered orientation tuning via a multiplicative gain enhancement without changing tuning width. Critically, endogenous attention enhanced sensitivity to SFs at, above and below the target SF, whereas exogenous attention preferentially enhanced SFs above the target SF. These findings provide evidence for the assumed differential SF profiles required to model the effects of these two types of attention on performance (Jigo et al., 2021).

## Materials and Methods

### Observers

The same six observers (aged 19-26; 4 females), including authors A.F. and S.O., participated in both experiments (15 experimental sessions per experiment). All observers had normal or corrected-to-normal vision and provided informed consent. All experimental procedures were in agreement with the Helsinki declaration and approved by the university committee on activities involving human subjects at New York University.

### Apparatus

Stimuli were generated using MATLAB (MathWorks, Natick, MA) and the Psychophysics toolbox (Brainard, 1997; Pelli, 1997; Kleiner et al., 2007). Observers sat in a dimly lit room with their head positioned on a chinrest 57 cm away from a 22-inch Iiyama Vision Master Pro 514-CRT monitor (1280 × 960 resolution; 100 Hz). A ColorCAL MKII Colorimeter was used for gamma correction (Cambridge Research Systems, Rochester, Kent, UK). Observers viewed the monitor display binocularly and fixation was monitored using an EyeLink 1000 eye tracker (SR Research, Ottawa, Ontario, Canada) at a sampling rate of 1000 Hz.

### Stimuli

Stimuli were presented on a gray background (luminance: 68 cd/m^2^) at four spatial locations denoted by placeholders composed of four black dots forming a square (3° wide; **Figure 1A**). The placeholders remained on the screen for the entire experimental session and were presented at each intercardinal location at 7° of eccentricity. The target was a random phase vertical Gabor generated by modulating a 2 cpd sine wave with a Gaussian envelope (0.8° SD). Four white noise patches were independently and randomly generated on each trial and presented inside each placeholder. The noise was bandpass-filtered to contain SFs between 1 and 4 cpd and scaled to have 0.12 root-mean-square contrast. On any given trial the test stimulus could be a noise patch or a target embedded in noise (**Figure 1A,B**). The same stimuli were used in both experiments.

**Figure 1.**
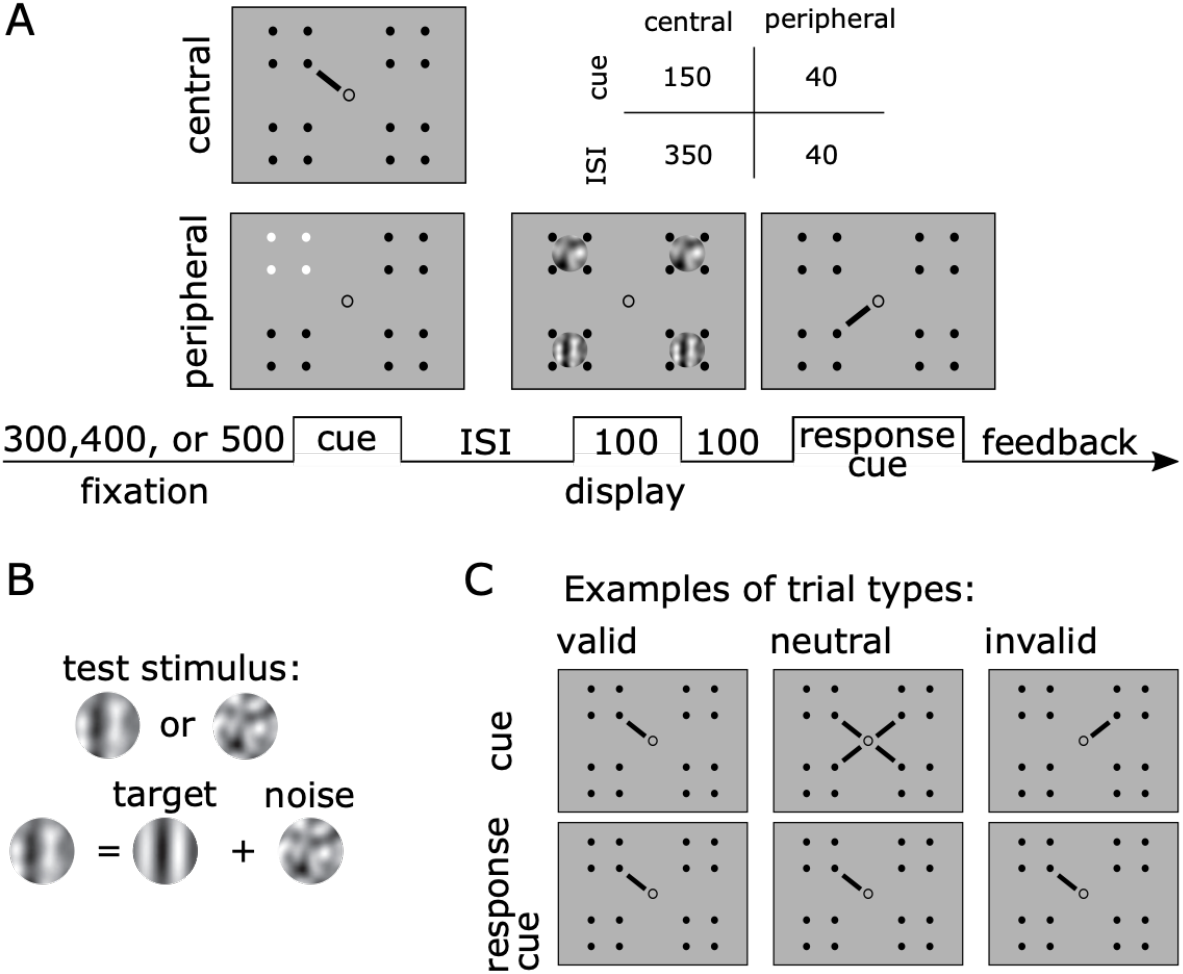
(**A**) *Trial Timeline in ms*. Observers performed two yes/no detection tasks. For endogenous attention (experiment 1), the cue was a line extending from the center indicating one spatial location (central cue). For exogenous attention (experiment 2), the placeholder at one spatial location briefly changed in polarity (peripheral cue). The timings (cue and ISI) differed for both experiments (central and peripheral cue) to optimize the effectiveness of the cues. (**B**) *Stimuli*. In half of the trials the test stimulus consisted of a Gabor embedded in noise, in the other half the stimulus was just noise. (**C**) *Trial types*. The example demonstrates valid, neutral, and invalid trial in Experiment 1. In Experiment 2, the cue and response cue mappings were the same, but the cue was peripheral instead of central.

### Experimental design and statistical analysis

#### Task Procedures

##### Experiment 1

Observers performed a visual detection task. A test stimulus, defined as the patch probed by the response cue, was presented at one of four possible spatial locations denoted by placeholders. In half of the trials, the test stimulus contained only noise; in the other half, the target was embedded in the noise. To diminish temporal expectation and ensure our attention effects were driven by the cues, each trial began with one of three equally likely fixation periods (300, 400, or 500 ms). After the fixation period, a pre-cue was presented for 150 ms, followed by a brief inter stimulus interval of 350 ms. Next, four patches (three irrelevant stimuli and one test stimulus), one inside each placeholder, were simultaneously presented for 100 ms. Each placeholder could encompass a patch composed of a target embedded in the noise or just noise with 50% probability. Thus, on any given trial, whether a target was presented was independently determined for all four locations.

The test stimulus was indicated by a central response cue, which was presented 100 ms after the offset of the patches and remained on the screen until response. The cue validity was manipulated such that 60% of the trials were valid, 20% were neutral, and the remaining 20% were invalid; from the cued trials indicating one location, 75% were valid trials (**Figure 1C**). Therefore, observers were incentivized to deploy their voluntary spatial attention to the location probed by the pre-cue as most of the trials were valid. In the valid and invalid trials, the pre-cue was presented as a central line pointing to one of the placeholders. In valid trials, the pre-cue matched the response cue; in invalid trials the pre-cue and response cue did not match. In neutral trials, all locations were pre-cued (4 lines, one pointing to each placeholder) to distribute attention to all placeholders. The observers’ task was to detect whether the test stimulus contained a target by pressing the “z” key for yes or “?” for no on an apple keyboard. On incorrect trials observers received auditory feedback in the form of a tone (400 Hz). At the end of each block observers also received feedback in the form of percent correct.

All attention conditions (valid, invalid, and neutral) were interleaved within each block. Observers completed 7 blocks for a total of 1008 trials per session. Each observer completed a training session (1008 trials) to familiarize themselves with the task. This was followed by 15 experimental sessions (15,120. During training, a contrast threshold—defined as the level of contrast needed to achieve 75% accuracy on neutral trials—was determined for the target using an adaptive titration method (Best PEST; Pentland, 1980). If an observer was under/over-performing by ≥5% on neutral trials on any given session, their contrast threshold was updated in the following session.

Stimulus presentation was contingent upon central fixation. Fixation was monitored from the start of the trial until response cue presentation. Trials with fixation breaks or blinks—defined as a deviation in eye-position by ≥1.5° from fixation—were repeated at the end of the block. On average observers 5.25% of trials in Experiment 1 and 1.58% of trials in Experiment 2 were repeated.

##### Experiment 2

The task was very similar to experiment 1 with slight differences to facilitate the effects of exogenous attention on performance. The attentional cues changed from central to peripheral (**Figure 1**) and were composed of a brief change in polarity of one (valid/invalid) or all (neutral) placeholder/s, and the duration of the cue and ISI was decreased to 40ms each (**Figure 1**). Cue validity was equated across valid, invalid and neutral trials (33% for each condition), making the cues uninformative.

#### Reverse correlation

Reverse correlation, assuming a general linear model approach (**Figure 2**; Ahumada 1996; Eckstein & Ahumada 2002; Wyart et al., 2012; Li et al., 2016; Fernández et al., 2019), was used to assess the computations underlying attention’s modulatory effect on performance. The noise image of the test stimulus was transformed from a space defined by pixel luminance intensities to a space defined by the contrast energy across orientation (−80 to 80 with 19 points equally spaced on a linear scale) and SF (from 1 cpd to 4 cpd with 15 points evenly spaced on a log scale) components in the noise. We took the noise images *S* across trials and convolved them with two Gabor filters *g* (same size as the target) in quadrature phase with the corresponding orientation *θ* and SF *f* to compute energy *E_θ,f_* for all components, which is given by the following expression:

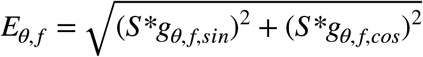

**Figure 2.**
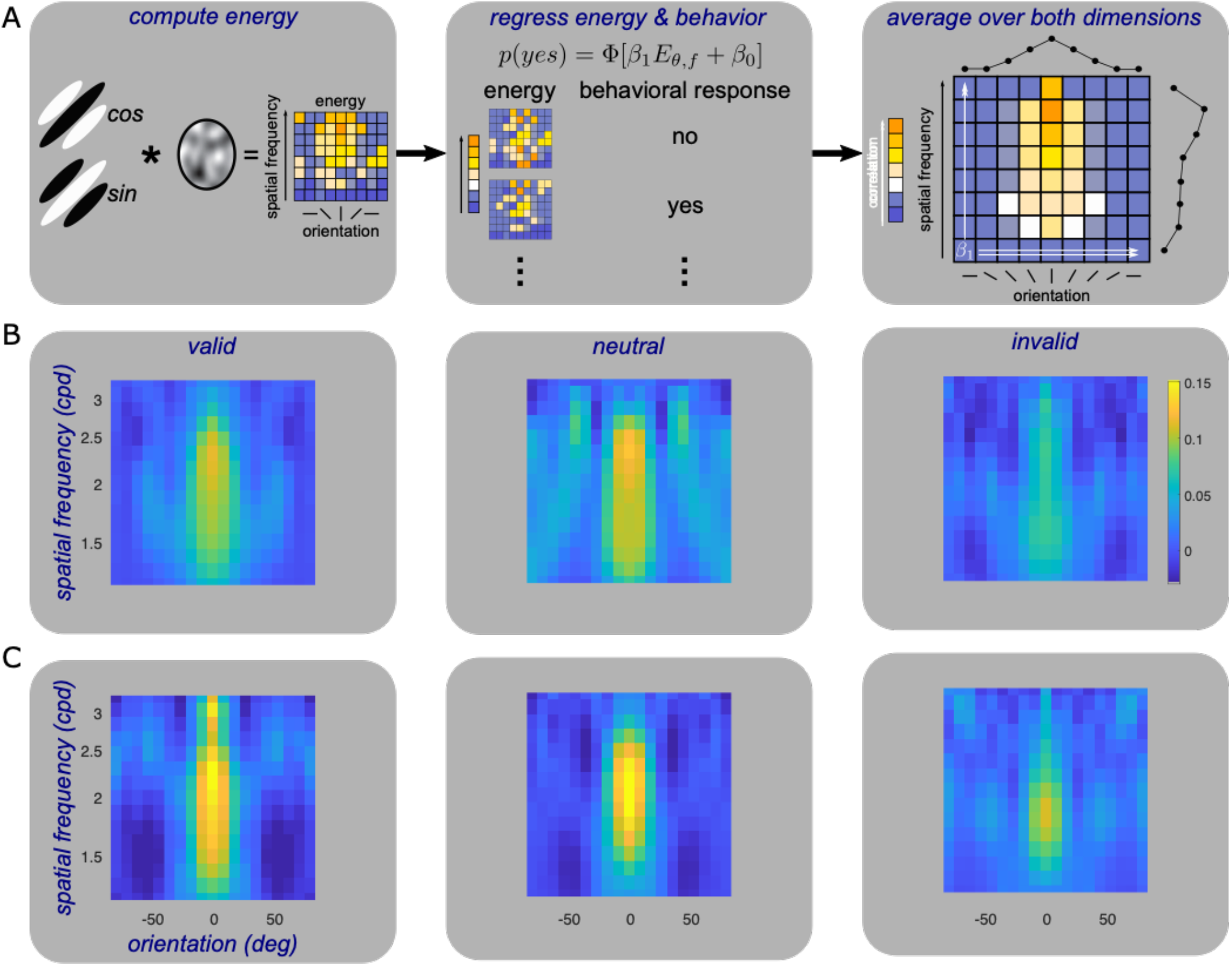
(A) *Reverse Correlation:* (1) Compute energy by convolving a bank of Gabor filters in quadrature phase with the noise patches; (2) Regress the trial-wise energy fluctuations with behavioral responses; (3) The slope of the regression indexes sensitivity; take the mean (marginals) of the slope across the orientation and SF dimension to compute tuning functions. (B) *Endogenous attention (Experiment 1) Kernels* for one individual observer. (C) *Exogenous attention (Experiment 2) Kernels* for another individual observer.

We regressed the energy of each component with behavioral responses using probit binomial regression. The regression model was expressed as such:

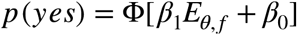

Where *β*_1_ is the slope of the regression and indicates the level of correlation between the energy and behavioral reports, *β*_0_ is an intercept term, Φ[.] indicates a probit link—inverse of the normal cumulative distribution function. We used the slope to index perceptual sensitivity. A slope of zero indicates that the energy of that component did not influence the observer’s responses. Prior to regressing, the energy of each component was sorted into a present or absent group and the mean of the energy for each group was subtracted and normalized to have a standard deviation of one, which allowed us to use only the energy fluctuations produced by the noise. The estimated sensitivity kernel *K* was a 2D matrix in which each element was a *β*_1_ value (**Figure 2A-C**). This process was completed independently for each of the three attention conditions.

SF and orientation tuning functions were computed by taking the marginals of the 2D sensitivity kernels. Marginals for orientation *θ_m_* were computed by taking the mean across the orientation dimension and were fit with a Gaussian of the form:

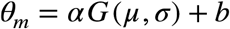

Where *α* controls the amplitude, *μ* the center, *σ* the width, and *b* the baseline or y-intercept. Conversely, marginals for SF *f_m_* were computed by taking the mean across the SF dimension and were fit with a truncated log parabola of the form (Watson & Ahumada, 2005):

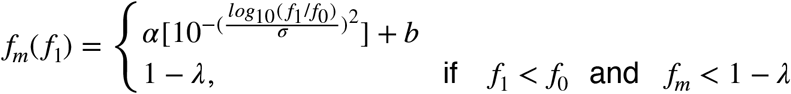

Where *f*_1_ are the tested SFs, *f*_0_ controls the peak frequency, *σ* controls the width, *λ* is a truncation parameter that flattens the curve, *α* scales the function, and *b* the baseline or y-intercept. The amplitude of the functions was computed by taking the difference between *α* and |*b*|. Endogenous attention was fit with a log parabola without truncation.

We conducted marginal reconstruction to determine whether we could treat orientation and SF as separate dimensions. We took the outer product of the marginal orientation and the marginal of the absolute value of the SF tuning functions to reconstruct the sensitivity kernels for all attention conditions. We then correlated the reconstructed sensitivity kernels with the original kernels. The higher the correlation the higher the separability.

#### Statistical analysis

A bootstrapping procedure was used to test for statistical significance across all fitted parameters (Li, Pan, Carrasco 2019, 2021). We first resampled, with replacement, each observers’ data to generate new 2D sensitivity kernels per the exogenous and endogenous attention conditions. We then averaged across observers to arrive at a new mean sensitivity kernel. This was followed by taking the marginals for orientation and SF and fitting them with a Gaussian and truncated log parabola, respectively. We then took the difference in fitted parameters between valid and invalid, valid and neutral, and neutral and invalid to generate difference scores. This process was repeated 1000 times to arrive at distributions of difference scores. Significance was defined as the proportion of the difference scores that fell above or below zero. When appropriate, e.g., when looking at the differences in the bootstrapped marginals, adjustments for multiple comparisons were conducted via Bonferroni correction (Holm 1979). When comparing between conditions we report 68% confidence intervals as it equals a standard error (Wichmann & Hill, 2001).

Parameters estimated from the bootstrapped data were also used to compare the attentional effects generated by endogenous and exogenous attention. First, parameter estimates across each bootstrapping iteration for the valid and invalid functions were subtracted to generate attentional effects. Second, we took the difference in attentional effects between endogenous and exogenous attention to generate difference-difference scores. Significance was defined as the proportion of the difference-difference scores that fell above or below zero. This process was repeated for benefits (valid-neutral) and costs (neutral-invalid).

Significance for detection sensitivity (*d*′), criterion (c), and reaction time was computed using within subjects ANOVAs and when necessary two-tailed *t*-tests. Effect sizes are reported as generalized eta squared (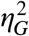; Bakeman, 2005) or Cohen’s *d* (Cohen, 1992). For all reported null effects, we provide a complementary Bayesian approach (Masson, 2011). The ANOVA sum of squared errors were transformed to estimate Bayes Factors *B* as well as Bayesian information criterion probabilities *pBIC* for the null *H*_0_ and alternative *H*_1_ hypothesis given the data *D*. We report Bayes Factors as odds in favor of the null hypothesis (e.g., *B*[4:1] indicates 4 to 1 odds in favor of the null hypothesis).

#### Model Comparisons

To assess which model best fit the orientation and SF tuning functions we conducted model comparisons using the Akaike information criterion corrected for small sample sizes (AICc). Assuming normally distributed error and constant variance, AICc can be computed from the estimated residuals (Burnham & Anderson, 2002):

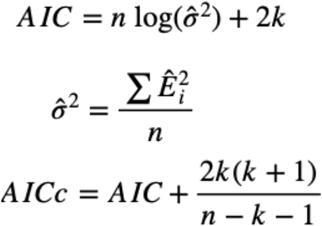

Here *n* is the number of data points, *k* is the number of parameters in the candidate model, and *Ê_i_* is the estimated residuals. We report differences in AICc scores (Δ*AICc*):

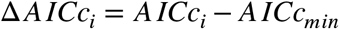

The best model is one where Δ*AICc_i_* ≡ Δ*_min_* ≡ 0. The level of support for model *i* is substantial if Δ*AICc_i_* falls within 0-2, considerably less if it falls within 4-7, and no support if >10 (Burnham & Anderson, 2002). We discarded all models with a Δ*AICc* > 10. In all cases the chosen model had a Δ*AICc* ⩽ 2.

Model comparisons were computed on the non-bootstrapped individual data. In the orientation dimension, eight models with different combinations of shared parameters for the valid, neutral, and invalid functions were tested (e.g. if only *σ* is shared then only one *σ* parameter was used to capture the width of all three cues, while three separate parameters for the gain (*α*) and baseline (*b*) were used for a total of seven [*k* = 7] free parameters). In the SF dimension, sixteen models were tested. For exogenous attention, the truncation term was always left free to vary across attentional cues. For endogenous attention, the truncation term was removed. Once the best model was determined, it was then fit to the bootstrapped data for statistical analysis.

## Results

### Experiment 1

We first examined the effect of endogenous attention on performance (**Figure 3**). Endogenous attention improved sensitivity (indexed by *d*′; F(2,10) = 42.88, *p* < 0.001, 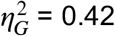) at the target location. Sensitivity for valid cue trials was higher than for the neutral (*t*(5) = 6.246, *p* = 0.002, *d* = 1.041), which in turn was higher than for invalid cues (t(5)=5.716, *p* = 0.002, *d*= 0.827). We also analyzed RTs as a secondary variable, to assess if there were any speed-accuracy tradeoffs. Attention sped responses (F(2,10) = 35.32, *p* < 0.001, 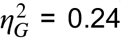). Observers were faster to respond when presented with a valid than a neutral cue (t(5)= −6.532, *p* = 0.001, *d* = −0.816), which in turn was faster than for an invalid cue (t(5)= −4.61, *p* = 0.005, *d* = 0.44). These results rule out any speed-accuracy tradeoffs. Furthermore, there were no significant differences in decision criterion across all attention conditions (F(2,10)= 2.006, *p* = 0.185; *B*[2.18:1]; *pBIC* (*H*_1_|*D*) = 0.314; *pBIC* (*H*_0_|*D*) = 0.686). In sum, sensitivity was improved at the attended location and impaired at the unattended location.

**Figure 3.**
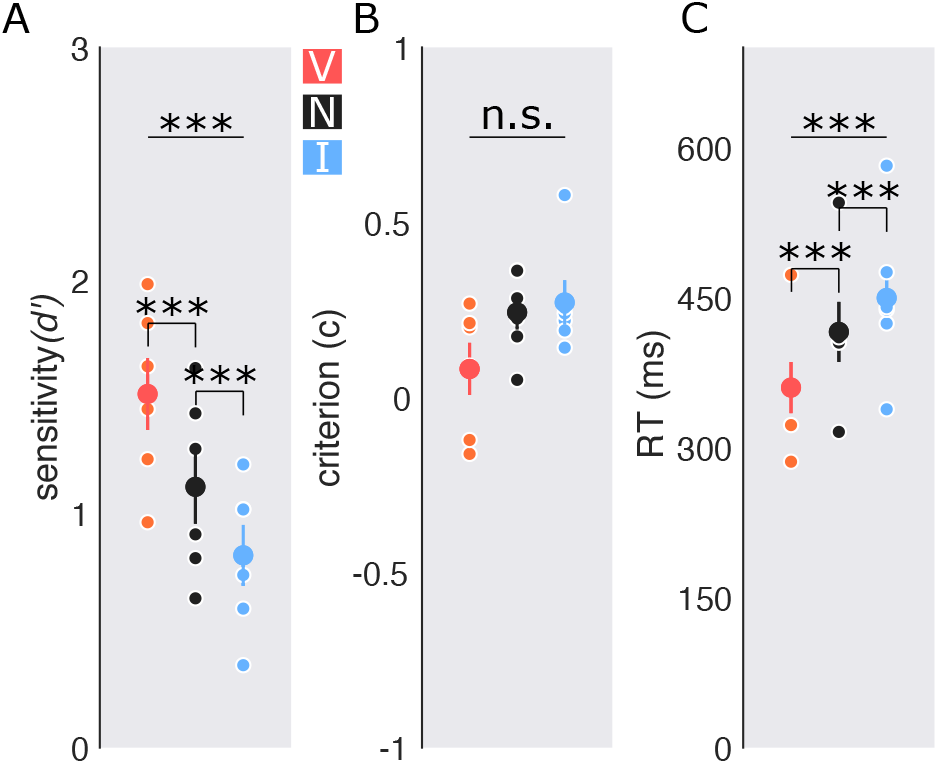
*Performance, Experiment 1: Endogenous attention*. (A) *Detection sensitivity indexed by d′ with central cues*. (B) *criterion*. (C) *Reaction time (RT) in milliseconds*. Big dots represent the group average; small dots= individual data. Error bars are ± 1 SEM. * *p* < 0.05; *** *p* < 0.001.

Having established that the cues successfully manipulated endogenous attention and affected performance, we next regressed the energy across trials with each observer’s responses using binomial regression. This provided a 2D sensitivity kernel where each component consisted of a *β*_1_ weight representing the level of correlation between reporting the target as present in the test stimulus and the energy in the noise. High weights indicate a high correlation, suggesting that this component in the noise positively influenced the observer’s responses, whereas weights close to zero or below indicate no correlation or a negative relation. As expected, the highest weights were observed at the components corresponding to the orientation and SF of the target.

To ensure that we could treat orientation and SF as separable dimensions in our data, we conducted a separability test via marginal reconstruction. We observed high levels of separability (0.90, 0.81, 0.89,0.87,0.80,0.93 for individual observers; mean: 0.87±0.05).

Having established that the feature dimensions are separable, we focused our analysis on the marginals of the 2D sensitivity kernels. We fit the orientation marginals with Gaussians and the SF marginals with truncated log parabolas.

Model comparisons revealed that all orientation tuning functions were best fit with a shared width parameter (*σ*) across attentional conditions; indicating that the width of the functions did not differ (**Figure 4A**). The valid central cue and the neutral cue increased the gain on orientations more than the invalid cue (*valid vs invalid*: *p* = 0.002; *neutral vs invalid*: *p* = 0.044; **Figure 4B-D**). Endogenous attention enhanced the gain across all orientations without changing tuning width. The best fit model in the SF dimension was also one with only a shared width parameter (*σ*) across all cues (**Figure 5A**). The valid central cue resulted in higher sensitivity—via an amplitude enhancement—than the invalid cue (*p*<0.001). A follow up bootstrapping analysis on the functions revealed that this enhancement spanned SFs below and above the target SF (1.43-2.32 cpd; Figure 5D-gray horizontal bar). Sensitivity for the valid and neutral cue did not significantly differ(*p*=0.45), but sensitivity for invalid cue was significantly lower than for the neutral cue (*p* < 0.001). Additionally, the valid cue shifted peak sensitivity to lower SFs than the neutral cue, *p*=0.018 (**Figure 5B-D**), and this was the case for 5 of the 6 individual observers (**Figure 5B**). Endogenous attention enhanced SFs at, above and below the target SF.

**Figure 5:**
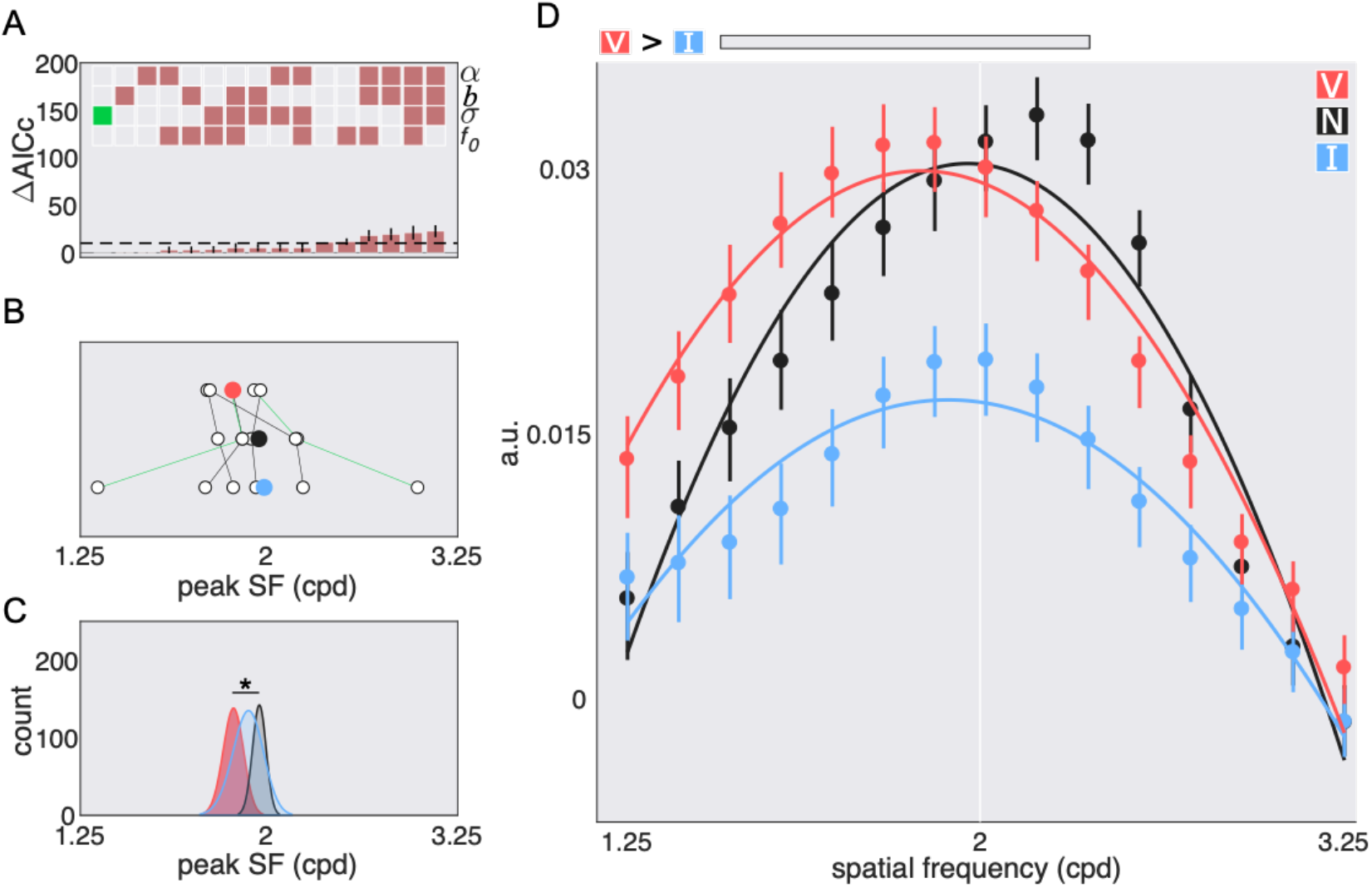
*SF tuning functions with central cues, Experiment 1: Endogenous attention*. (A) model comparison results: gain *α*, width *σ*, baseline *b*, and peak *f*_0_; empty squares = not shared; filled in squares = shared; green square = chosen model. Dashed line = cutoff (10). (B) Parameter estimates for the peak of the fit to the *individual* data of the chosen model; colored circles=mean; green lines=authors’ data. (C) Parameter estimates for the peak of the fit to the *bootstrapped* data of the chosen model. (D) Best fit tuning functions (using the chosen model) to the group averaged data. Gray horizontal bars = significant differences after correction for multiple comparisons; the units are arbitrary (a.u.). Error bars are 68% C.I. * *p* < 0.05. ** *p* < 0.01, * *p* < 0.05

### Experiment 2

Exogenous attention also improved performance (**Figure 6**). Peripheral cues enhanced detection sensitivity at the target location (F(2,10)=17.06, *p* < 0.001, 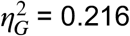); sensitivity was higher for the valid than the neutral cue (*t*(5)=4.159, *p* < 0.001, *d*= 0.649), which in turn was higher than invalid cues (t(5)=3.8, *p* < 0.012, *d* = 0.533). The type of cue also modulated reaction time (F(2,10)=7.299, *p* = 0.011, 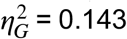). Responses were faster for valid than neutral (*t*(5)=−2.918, *p* = 0.033, *d*= −0.610) and invalid (*t*(5)=−2.707, *p* = 0.042, *d*= −0.875), and marginally faster for neutral than invalid cues(*t*(5)=−2.175, *p* = 0.082). There were no speed-accuracy tradeoffs. Decision criteria did not differ across attention conditions (F(2,10)<1; *B*[4.55:1]; *pBIC*(*H*_1_|*D*) = 0.179;*pBIC*(*H*_0_|*D*) = 0.821). In sum, sensitivity was improved at the attended location and impaired at the unattended location.

**Figure 6.**
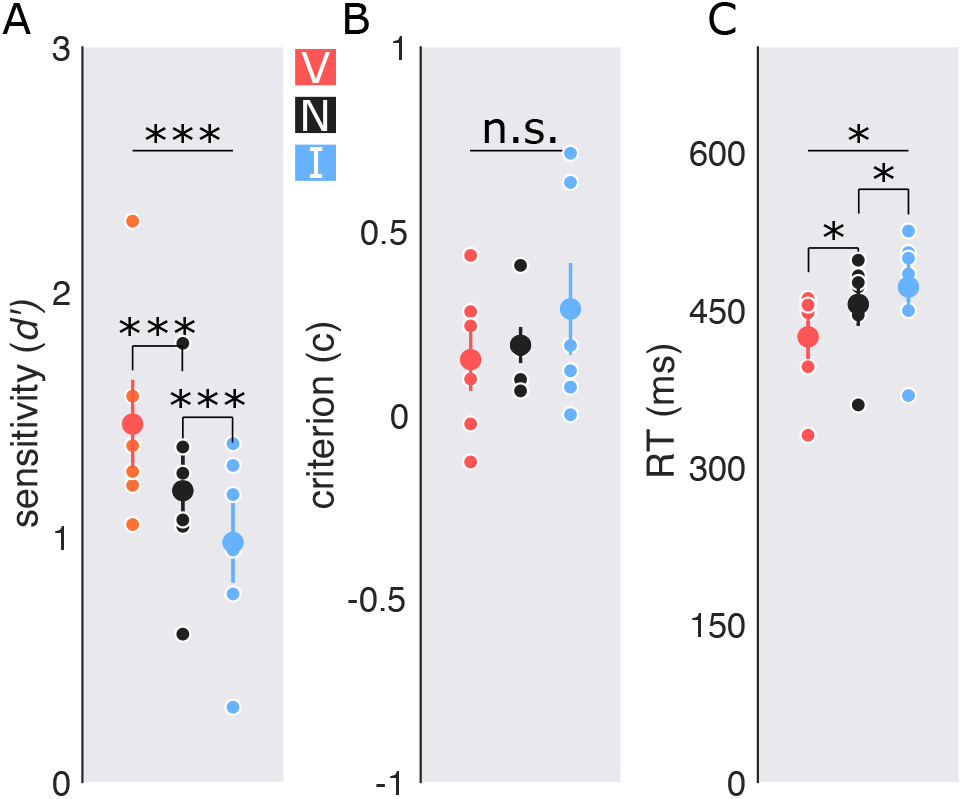
*Performance, Experiment 2: Exogenous attention*. (A) *Detection sensitivity indexed by d’ with peripheral cues*. (B) *criterion*. (C) *Reaction time (RT) in milliseconds*. Big dots represent the group average; small dots= individual data. Error bars are ± 1 SEM. * *p* < 0.05; *** *p* < 0.001.

When examining the 2D sensitivity kernels, much like in Experiment 1, the highest weights also corresponded to the components representing the orientation and SF of the target and there was a high level of separability between both feature dimensions (0.91, 0.79, 0.77, 0.77, 0.82, 0.81 for individual observers; mean: 0.81±0.05).

To facilitate comparisons with endogenous attention we fit the orientation data with only a common width (*σ*) parameter (*2^nd^ best model:* Δ*AICc* = 2; **Figure 7A**). The difference between the two best models was negligible as the level of support for Δ*AICc* differences of 0-2 is the same (Burnham & Anderson 2002). The valid peripheral cue enhanced the gain (*α*) across all orientations more so than the neutral, *p* < 0.001, or invalid cue, *p* < 0.001 (**Figure 7B-D**). Exogenous attention altered orientation representations by boosting the gain without changing tuning width.

**Figure 7:**
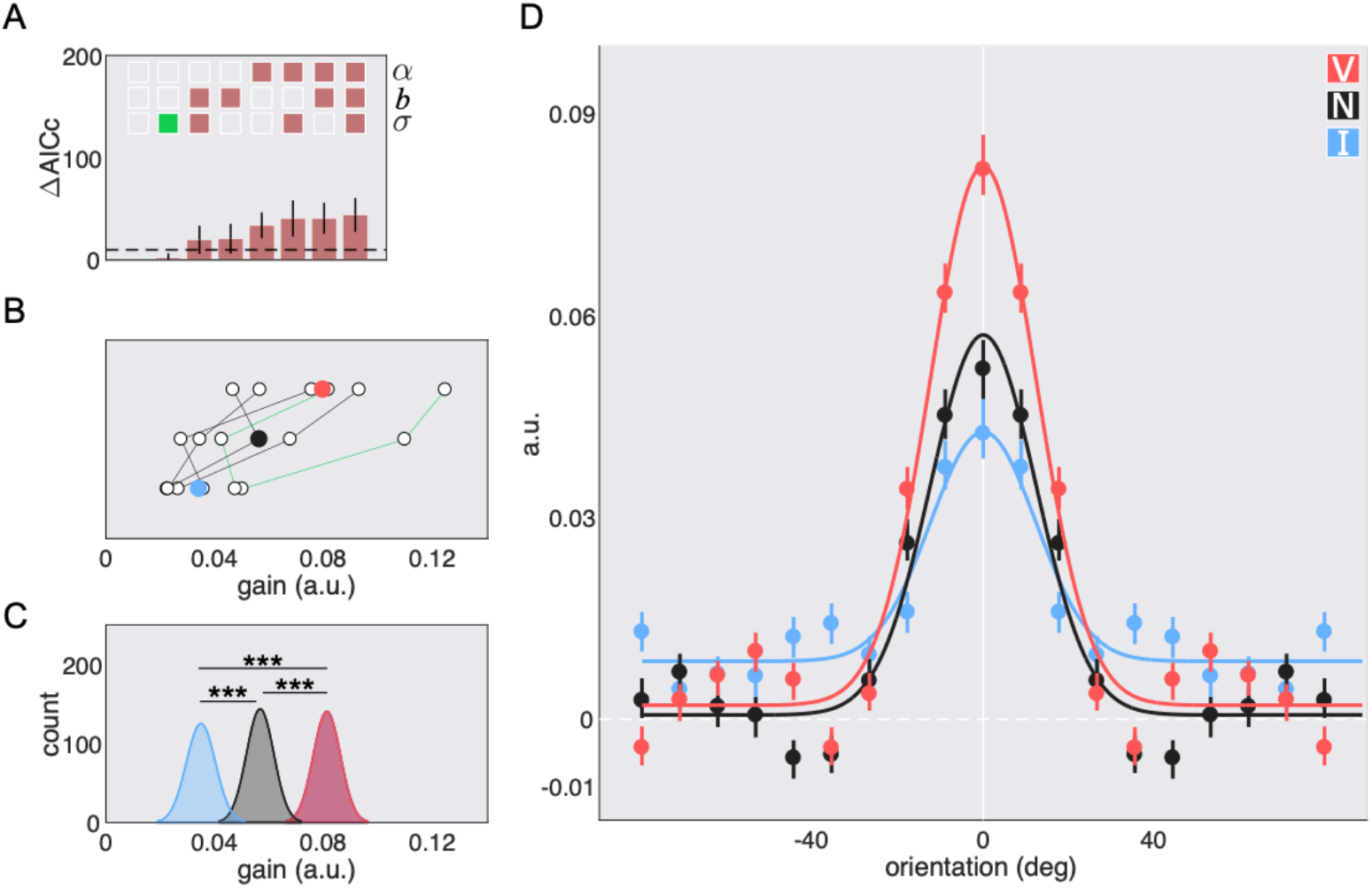
*Orientation tuning functions with peripheral cues, Experiment 2: Exogenous attention*. (A) model comparison results: gain *α*, width *σ*, and baseline *b*; empty squares = not shared; filled in squares = shared; green square = chosen model. Dashed line = cutoff for significance. (B) Parameter estimates for the gain of the fit to the *individual* data of the chosen model; colored circles=mean; green lines=authors’ data. (C) Parameter estimates for the gain of the fit to the *bootstrapped* data of the chosen model. (D) Best fit tuning functions (using the chosen model) to the group averaged data; the units are arbitrary (a.u). Error bars are 68% C.I. *** *p* < 0.001.

In the SF domain, the best fit model also contained only a shared parameter for the width (**Figure 8A**). The valid cue shifted peak sensitivity (*f*_0_) to higher SFs when compared to the neutral (*p* = 0.011) and invalid cues (*p* = 0.003). Additionally, the valid function had greater amplitude than the neutral (*p* = 0.003) and invalid functions (*p*=0.04). Further examination of the bootstrapped marginals revealed that this difference was driven by SFs above the target SF (valid vs neutral: 2.16-2.84 cpd; valid vs invalid: 2.31-2.64 cpd; **Figure 8B-D**). In the individual data (**Figure 8B**), for all six observers the SF peak for the valid functions is clearly above the SF peak for the invalid and for five the SF peak for the valid is above the peak of neutral. No differences in the amplitude of the valid, neutral, and invalid functions for SFs below the target SF survived corrections for multiple comparisons. Exogenous attention preferentially enhanced sensitivity to SFs higher than the target SF.

**Figure 8:**
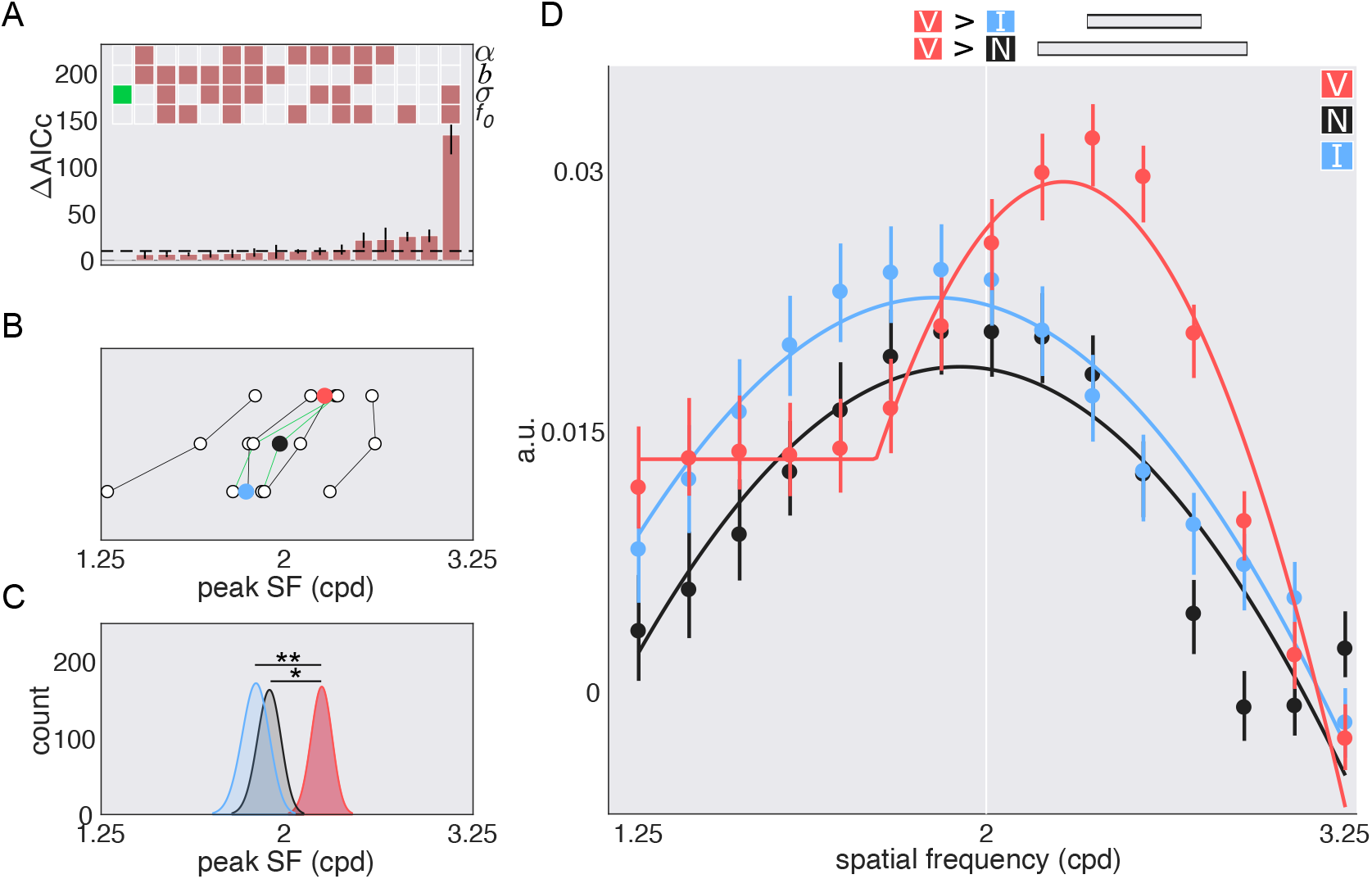
*SF tuning functions with peripheral cues, Experiment 2: Exogenous attention*. (A) model comparison results: gain *α*, width *σ*, baseline *b*, and peak *f*_0_; empty squares = not shared; filled in squares = shared; green square = chosen model. Dashed line = cutoff (10). (B) Parameter estimates for the peak of the fit to the *individual* data of the chosen model; colored circles=mean; green lines=authors’ data. (C) Parameter estimates for the peak of the fit to the *bootstrapped* data of the chosen model. (D) Best fit tuning functions (using the chosen model) to the group averaged data. Gray horizontal bars = significant differences after correction for multiple comparisons; the units are arbitrary (a.u.). Error bars are 68% C.I. * *p* < 0.05; ** p < 0.01

### Comparisons—endogenous vs. exogenous attention

#### Effects on detection sensitivity

To ensure that differences between endogenous and exogenous attention in sensory tuning are not driven by changes in detection sensitivity; we assessed whether both endogenous and exogenous attention affected task performance in a similar fashion. Both types of covert attention similarly affected detection sensitivity (*d*′; F(1,5)<1; *B*[3.94:1]; *pBIC* (*H*_1_|*D*) = 0.202; *pBIC*(*H*_0_|*D*) = 0.798) and the effect of the cues did not depend on the type of attention being deployed (F(2,10)=2.77; *p*=0.109; *B*[1.58:1]; *pBIC*(*H*_1_|*D*) = 0.387; *pBIC*(*H*_0_|*D*) = 0.613).

#### Sensory tuning

To compare whether and how endogenous and exogenous attention differentially alter sensory representations we examined the difference in overall attentional effects (valid-invalid), ‘benefits’ (valid-neutral), and ‘costs’ (neutral-invalid) indexed by differences in parameter estimates (**Figure 9**). For attentional effects, exogenous attention exhibited a greater difference in orientation gain between the valid and invalid functions than endogenous attention (**Figure 9A**; *p* < 0.001). However, the opposite was true in the SF dimension (**Figure 9A**; *p* < 0.001). Additionally, peak SF was higher for the valid than invalid functions with exogenous but not in endogenous attention (**Figure 9A**; *p* < 0.001). In both the orientation and SF dimensions the gain enhancements by exogenous attention were greater than by endogenous attention (**Figure 9B**; all *p*’s < 0.001). Exogenous valid cues shifted peak SF above the neutral peak but endogenous attention did not (**Figure 9B**; *p* < 0.001). Additionally, the invalid cue impaired orientation gain to a larger extent with exogenous than endogenous attention (**Figure 9C**; *p* < 0.001). In contrast, invalid cues impaired SF amplitude more with endogenous than exogenous attention (**Figure 9C**; *p* < 0.001). Lastly, SF peak between the neutral and invalid functions between both types of attention was only marginally significant (*p* = 0.075), with the effect being more pronounced for exogenous than endogenous attention.

**Figure 9:**
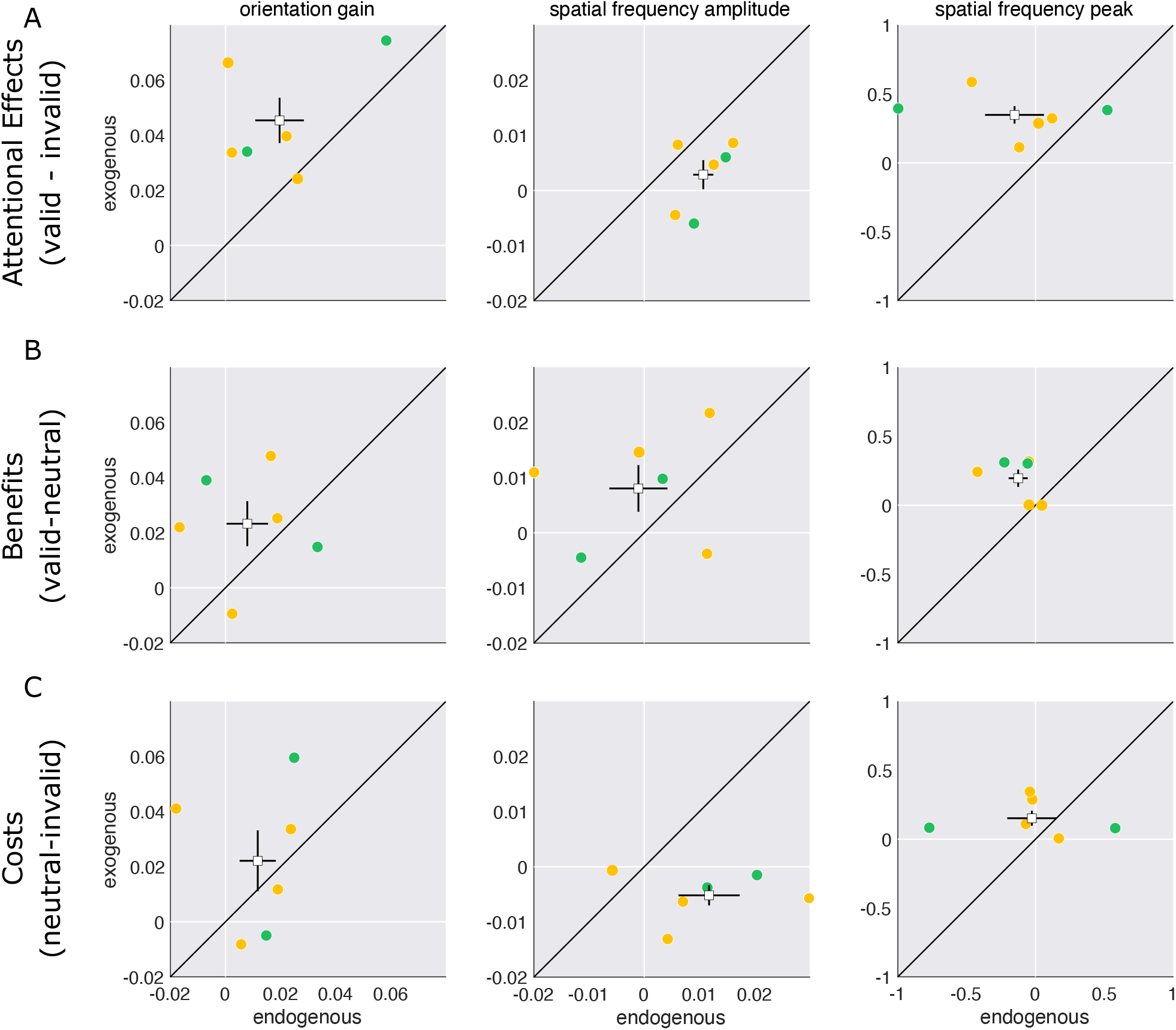
*Attentional Effects, Benefits & Costs*. Scatter plots of attentional effects (A; valid – invalid), benefits (B; valid-neutral) and costs (C; neutral-invalid) for endogenous vs. exogenous attention computed from fits to the individual data from parameter estimates for *orientation gain, SF amplitude*, and *SF peak*. orange circles=individual observers; green circles=author’s data; white squares = group means. Error bars are ±1 SEM.

## Discussion

We have compared the effects of endogenous and exogenous attention on orientation and SF tuning, with the same observers, task and stimuli. Both endogenous and exogenous attention yielded similar benefits in detection sensitivity at the attended location and costs at unattended locations, ruling out performance differences as a possible explanation for the differences in sensory tuning. In both experiments, reaction times were fastest for valid and slowest for invalid cues, ruling out speed-accuracy trade-offs, and there were no differences in decision criterion amongst the valid, neutral, and invalid cues.

### Orientation tuning

For both endogenous and exogenous attention, we found only gain changes, consistent with previous studies (endo: Wyart et al., 2012; Paltoglou & Neri, 2012; Barbot et al., 2014; exo: Baldassi & Verghese, 2005; Fernández et al., 2019).

Comparing overall attentional effects as well as the magnitude of benefits and costs between endogenous and exogenous attention revealed a stronger modulation by exogenous than endogenous attention. A possible explanation relates to the involuntary vs. voluntary nature of the effect; exogenous attention exerts its effects automatically to the same degree regardless of cue validity whereas the effect of endogenous attention increases with cue validity (Giordano et al., 2009). Although cue validity for endogenous was high (75%), the effects could have been more pronounced had the cue been valid 100% of the time. This gain difference could also be related to attentional modulations of neural activity by exogenous attention being approximately constant from early (V1,V2,V3: Müller & Kleinschmidt, 2007; Müller & Ebeling, 2008; Dugué et al., 2020) to intermediate (V3A,hV4,LO1: Dugué et al., 2020) visual areas, but increasing from early to intermediate visual areas (Dugué et al., 2020; MT, VIP: Maunsell & Cook, 2002) to parietal and frontal areas (Kastner et al., 1999) with endogenous attention. Consistent with the idea that endogenous attention is a top-down modulation from frontal and parietal areas feeding back to visual cortex (reviews: Chica et al., 2013; Beck & Kastner, 2009). In any case, future research could assess whether this result is due to a mechanistic effect of attention or to the experimental parameters used here.

### SF tuning

The SF data for both experiments was fit with the same model—shared parameter for the width. For endogenous attention, there was an overall increased amplitude of SFs above and below the target SF; this amplitude difference was primarily driven by the invalid cue. The overall amplitude difference is consistent with endogenous attention excluding external noise without changing SF tuning (Lu & Dosher, 2004). Additionally, peak sensitivity shifted to lower SFs than the neutral curve. Endogenous attention could have flexibly shifted sensitivity to lower SFs to counteract any effect of masking by the target SF (Meese & Hess, 2004). Exogenous attention shifted peak sensitivity to SFs higher than those for the neutral and invalid functions, which did not differ between themselves. This valid shift led to a preferential gain enhancement at SFs above the target SF.

Psychophysical studies using both reverse correlation (Fernández et al., 2019) and critical-band-masking (Talgar et al., 2004) found no change in SF tuning with exogenous attention. Here, we capitalized on previous and recent psychophysical findings to improve our task design. We hypothesize that the biggest contributors for the observed preferential enhancement of high SFs with exogenous attention was: (1) placing stimuli at less eccentric locations and (2) increasing the target SF. In our previous study (Fernández et al., 2019), the stimuli were placed at 10° eccentricity, and the target SF was 1.5cpd, outside the optimal range of cueing benefits with exogenous attention (Jigo & Carrasco, 2020). By decreasing the eccentricity and increasing the target SF, we optimized stimulus parameters and could observe shifts in sensory tuning induced by exogenous attention.

A recent study has also revealed differential effects of exogenous and endogenous attention on contrast sensitivity (indexed by *d*′; Jigo & Carrasco, 2020). Exogenous attention preferentially enhances SFs higher than the intrinsic peak frequency in a neutral baseline condition, whereas endogenous attention provides similar benefits at a broad range of SFs above and below the peak at baseline. Conventional signal detection theory is agnostic as to the underlying computation. Here, using reverse correlation, we show the computation (gain changes) that may underlie the differential effects of endogenous and exogenous attention and compare differences in gain modulation between both types of attention.

Reverse correlation typically produces tuning curves that are similar to those derived from neural recordings (Neri & Levi, 2006). At the heart of reverse correlation is linking task properties (e.g., stimuli) to responses (behavioral or neural). Given the current findings, future electrophysiological studies focused on primate visual cortex, could assess whether gain modulations by endogenous attention would be similar for neurons coding SFs lower and higher than those of the target’s SF, whereas gain modulations by exogenous attention would be asymmetrical, preferentially enhancing the gain of neurons tuned to higher SFs than the target’s SF.

Normalization is a canonical neural computation (Carandini & Heeger, 2012) in both human (Bloem & Ling, 2019) and non-human (Ni & Maunsell, 2017, 2019) primates. A prominent normalization model of attention links attentional gain changes reported in electrophysiology and human psychophysics (Reynolds & Heeger 2009). Recently, this normalization framework has been extended to model how endogenous and exogenous attention differentially alter visual perception (Jigo et al., 2021). To capture the effects of attention on contrast sensitivity (e.g., Jigo & Carrasco, 2020), texture segmentation (e.g., Yeshurun & Carrasco, 1998; Barbot & Carrasco, 2017) and acuity (e.g., Montagna et al., 2009) the model requires SF tuning profiles consistent with the ones reported here: (1) a narrow high SF enhancement with exogenous attention; (2) a broad SF profile with endogenous attention. Indeed, without a differential SF profile the model cannot capture these established behavioral differences. Critically, these SF profiles scale the stimulus drive (i.e., bottom-up drive from simulated complex cells in V1), suggesting that these differential effects between endogenous and exogenous attention arise at a sensory rather than decision stage.

Our results can inform neurobiologists as human and non-human primates have similar spatial vision. For example, sensitivity peaks around the fovea and declines as eccentricity increases (Sasaoka et al., 2005), and sensitivity peaks in the middle of the SF range (2-5 cpd) and drops off for lower and higher frequencies (Harwerth & Smith, 1985). Similarities also extend to attentional modulations. For instance, attention improves contrast sensitivity and acuity in human (Herrmann et al., 2010; Montagna et al., 2009) and non-human primates (Pham et al., 2018; Golla et al., 2004).

What is the functional significance of the observed differences in SF tuning between endogenous and exogenous attention? For circumstances in which increasing resolution is not beneficial (e.g., driving through fog) the flexible, voluntarily controlled endogenous attention would be necessary. In fight or flight situations, an exogenous attentional system that is quick, reactive, and inflexibly increases resolution would be beneficial. Mice exhibit primate-like behavioral signatures of both endogenous and exogenous attention (You & Mysore, 2020) despite having poorer contrast sensitivity (Histed et al., 2012) and no fovea. Additionally, mice show increased gain in neurons tuned for high SFs under a state of heightened attention, thereby increasing their spatial acuity (Mineault et al., 2016). Despite pronounced differences in the visual system of mice and primates, a similar mechanism (e.g., attention) may underlie sensory tuning properties in mammals. Therefore, understanding how attention reshapes the sensory tuning of basic visual dimensions is of critical importance. Furthermore, our findings help constrain models of spatial vision and of visual attention by furthering our knowledge of the neural computations underlying the effects of attention on basic dimensions of spatial vision. Specifically, they provide evidence for the assumed differential SF profiles required to model the effects of these two types of attention on performance (Jigo et al., 2021).

To conclude, this study reveals the differential effects of endogenous and exogenous attention on sensory tuning. Both types of covert attention modulate orientation at the attended location by boosting the gain of all orientation channels without changing tuning width. This boost was greater with exogenous than endogenous attention. In the SF domain, endogenous attention enhanced the gain of SFs above and below the target SF whereas exogenous attention only enhanced those above. We propose these changes in sensory tuning may underlie the differential effects of endogenous and exogenous attention on performance.

The authors have no conflict of interest to declare.

## Acknowledgements

This research was supported by the National Eye Institute of the National Institutes of Health under award number R01 EY019693 to M.C. and the National Institute of Neurological Disorders and Stroke of the National Institutes of Health under award number F99NS120705 to A.F. We thank Marc Himmelberg, Michael Jigo and Hsin-Hung Li, as well as other Carrasco lab members, for helpful comments.

